# Assessing Heterogeneity in the N-Telopeptides of Type I Collagen by Mass Spectrometry

**DOI:** 10.1101/2024.03.31.587441

**Authors:** Zsuzsanna Darula, Maxwell C. McCabe, Alex Barrett, Lauren R. Schmitt, Mark D. Maslanka, Anthony J Saviola, Joseph Orgel, Alma Burlingame, Claudia A. Staab-Weijnitz, Kurt Stenmark, Valerie Weaver, Robert J. Chalkley, Kirk C. Hansen

## Abstract

Collagen cross-links created by the lysyl oxidase and lysyl hydroxylase families of enzymes are a significant contributing factor to the biomechanical strength and rigidity of tissues, which in turn influence cell signaling and ultimately cell phenotype. In the clinic, the proteolytically liberated N-terminal cross-linked peptide of collagen I (NTX) is used as a biomarker of bone and connective tissue turnover, which is altered in several disease processes. Despite the clinical utility of these collagen breakdown products, the majority of the cross-linked peptide species have not been identified in proteomic datasets. Here we evaluate several parameters for the preparation and identification of these peptides from the collagen I-rich Achilles tendon. Our refined approach involving chemical digestion for protein solubilization coupled with mass spectrometry allows for the identification of the NTX cross-links in a range of modification states. Based on the specificity of the enzymatic cross-linking reaction we utilized follow-up variable modification searches to facilitate identification with a wider range of analytical workflows. We then applied a spectral library approach to identify differences in collagen cross-links in bovine pulmonary hypertension. The presented method offers unique opportunities to understand extracellular matrix remodeling events in development, aging, wound healing, and fibrotic disease that modulate collagen architecture through lysyl-hydroxylase and lysyl-oxidase enzymes.

## Introduction

A relatively under-explored, yet prominent post-translational modification (PTM) is an enzymatic modification of lysine by lysyl hydroxylase (LH) and lysyl oxidase (LOX) enzymes to generate protein-protein cross-links. The collagen family of proteins, the most abundant proteins in the human body^1,2^, represent a basic building block within almost every tissue and organ. Collagen I is found bundled into collagen fibers that exhibit a regular alternating pattern (D-banding) based on a ∼1/5 stacking offset (Figure 1A). The fibers are stabilized by a series of PTMs facilitated by the LH and LOX family of enzymes. Prior to triple helix formation, the LH enzyme family hydroxylates lysine (Lys) residues in the endoplasmic reticulum (ER), generating hydroxylysine (Hyl). Both Lys and Hyl are substrates of the extracellular matrix (ECM) cross-linking enzymes (LOX & LOXL1-4) which, via a lysine tyrosylquinone cofactor, catalyze the conversion of Lys or Hyl residues to δ-semialdehydes, allysine (alLys) or alhydroxylysine (alHyl), respectively (Figure 1B), in the extracellular space^3^. Dependent on whether Lys or Hyl act as precursors, different divalent and trivalent crosslinks are generated (Figure 1B, Supplemental Figure 1): The alLys aldehyde further reacts with a proximal Lys to form a divalent cross-link dehydro-lysinonorleucine (deH-LNL) via Schiff base formation, or with a hydroxylysine (Hyl) to form dehydro-hydroxylysinonorleucine (deH-HLNL) that can convert into lysino-5-ketonorleucine (LKNL). These two cross-links are reduced to lysinonorleucine (LNL) and 5- hydroxylysylnorleucine (HLNL), respectively. Activation of Hyl by a LOX family member, and further reaction with a neighboring Hyl generates dehydro-dihydroxylysinonorleucine (deH-DHLNL) which converts to hydroxylysino-5-ketonorleucine (HLKNL) that is reduced to 5,5’ dihydroxylysinonorleucine (DHLNL) (Figure 1B). Likewise, reaction with another alLys forms an aldol-product that reacts with a Lys or Hyl to form trivalent cross-link pyrroles or pyridinolines (hydroxylysylpyridinoline), otherwise known as pyridinoline (PYD), (Figure 1C) and deoxypyridinoline (DPD), or another aldol product to make tetravalent desmosines. Iso/desmosine species are reported to be exclusive to elastin cross-linking^4^. Several excellent reviews are available for further information about this class of enzymes and the resulting cross-linked species^5–10^.

**Figure 1.**
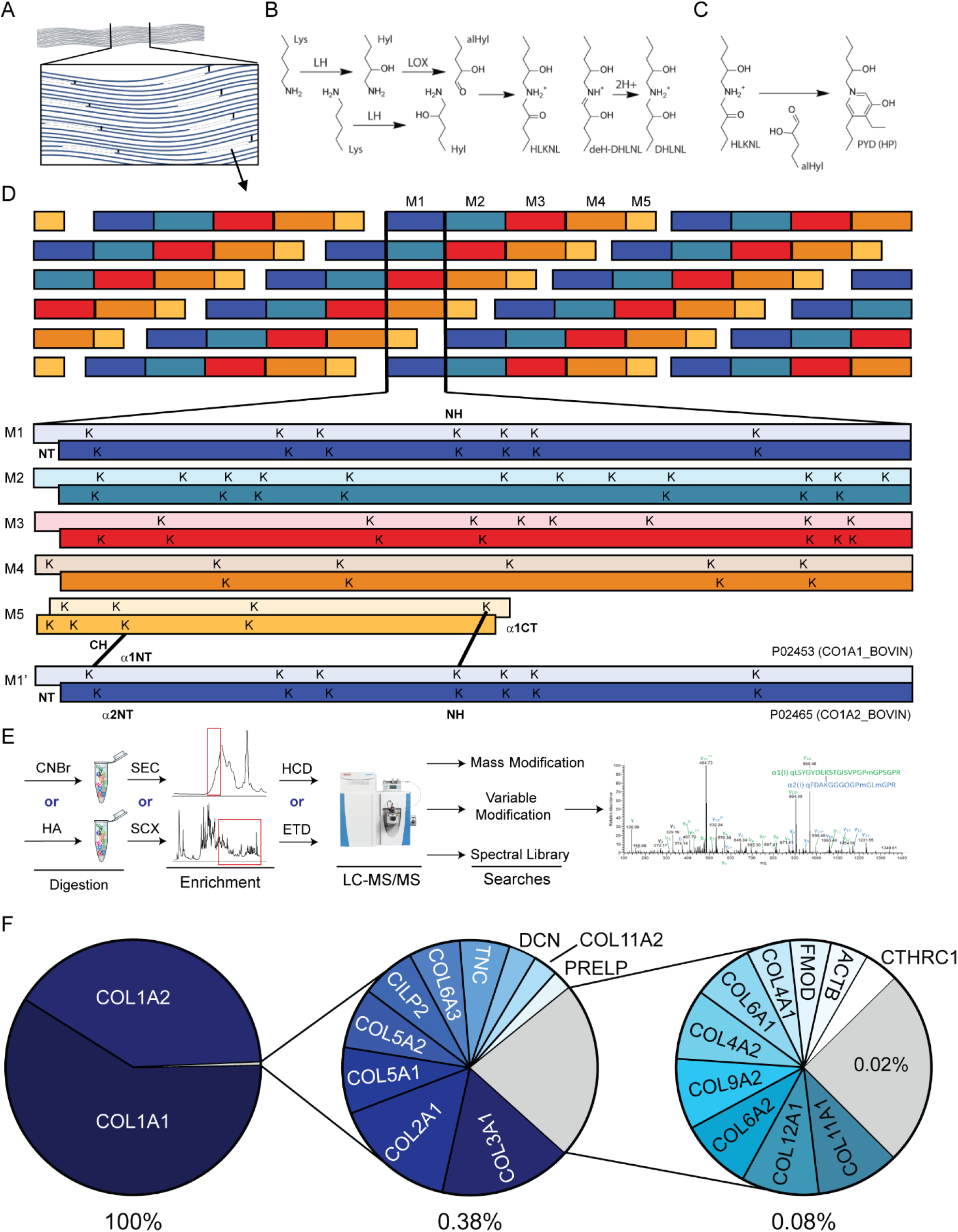
Collagen cross-linking and proteomic approach used for NTX peptide characterization. A) Overview of collagen fiber organization. B) LH and LOX modification of lysines followed by spontaneous cross-linking to form HLKNL product. Reduction to DHLNL is also shown. C.) Generation of tri-functional PYD cross-links D.) D-banding pattern observed in collagen fibers and corresponding lysine alignment across the 5 registries. Reported sites for COL1 cross-linking are labeled in bold: N-terminal (NT), C-terminal (CT), N-helix (NH), C-helix (CH). N-telopeptides from M1’ are in alignment with the M5 COL1A1 site K946 and COL1A2 K1021. E.) Analytical approach used to provide multiple lines of evidence for identified NTX species (further detail found in Supp. Figure 2). F.) Overall composition of the bovine Achilles tendon sample used for method development.

LOX enzymes have been shown to play a role in the generation of pathologic tissue microenvironments in the progression of several diseases^11–16^. In animal models, LOX enzymes have been shown to limit the reversal of fibrotic disease^17,18^, promote tumor progression^19–21^, and cause preconditioning of distant organs before metastatic dissemination^22,23^. In addition, a strong relationship has been shown between cross-linked products and tissue stiffness^14,19^ which results in intracellular signaling and activation of specific transcriptional programs^24^. However, very little is known about the substrate specificity of the LOX family members or the resulting architectures of modified collagens. This is, in part, due to challenges associated with the identification of cross- linked peptide species from complex tissue samples.

The best-known substrate of LOX is collagen I, and the cross-linked N-telopeptide (NTX) and C-telopeptide (CTX) of this protein have been used as clinical markers for several decades^25–27^. These assays are antibody-based, usually produced from clones generated against the linear peptides, such as QLSYGYDEKSTGGISVP^27^. It is not likely that these antibodies are specific for a single molecular cross-link, nor is it likely that the antibody has affinity for all possible forms of NTX cross-linked species.

The combination of offline fractionation and mass spectrometry (MS) has been used to identify cross-links from bovine, rat, and mouse collagens^8,28,29^. For example, previous work characterizing collagen I cross-links utilizing offline fractionation with high-resolution MS identified N- and C-terminal telo cross-linked species from bone^30^. The Eyre lab has utilized MS to map cross-links in bone and cartilage over the past four decades^8,29,31,32^ providing extensive knowledge regarding collagen cross-linking in several tissues and biological contexts. In addition, tandem MS methods identified an S-hydroxylysyl-methionine cross-link in the non-collagenous domain (NC1) of collagen IV hexamers^33,34^ which had previously been reported in the crystal structure^35^. These efforts, while informative, have relied on extensive, yet rare, domain expertise and almost exclusively utilized manual interpretation and identification strategies.

In the present study, we provide an analytical workflow that is compatible with modern proteomic approaches and database search routines that can be used for the identification of NTX cross-links. Several analytical variables were evaluated for COL1 NTX identification. Finally, we applied variable modification searching and spectral library methods to identify and facilitate the quantification of cross-link alterations in arteries from a large mammal model of hypoxia-induced pulmonary hypertension (hPH). The identified NTX peptides can be targeted for future biochemical and clinical studies to better understand the relationship between the LH and LOX enzymes, their activities, related disease states, interventions, and ultimately clinical outcomes.

### Materials and Methods Tissue Extraction

#### Bovine Ligament

Bovine Achilles tendon was obtained from Elastin Products (Product #C206). Approximately 10 mg of dry tissue was digested overnight with cyanogen bromide (CNBr) or hydroxylamine (HA) as previously described^36,37^. The digests were then treated with trypsin (1:50 enzyme:substrate ratio) using the FASP method^38^ using 10 kDa molecular weight cutoff filters (Sartorius Vivacon 500 #VN01H02) and chromatographically fractionated.

#### Bovine Main Pulmonary Artery

Main pulmonary artery (mPA) samples were extracted as previously described^37^. Briefly, approximately 4-5 mg of sample was decellularized using 3 washes of high salt/CHAPS buffer (50 mM Tris-HCl (pH 7.4), 0.25% CHAPS, 25 mM EDTA, 3 M NaCl) with agitation during each wash. Decellularized samples were subjected to extraction with 6M guanidine hydrochloride (Gnd-HCl), 100 mM ABC, pH 9.0 to remove chaotrope-soluble ECM components (soluble ECM), followed by chemical digestion using HA to solubilize fibrillar and other insoluble ECM components (insoluble ECM). Proteolytic digestion of the insoluble HA extract was carried out according to the filter-aided sample preparation (FASP) protocol^38^ with 10 kDa molecular weight cutoff filters (Sartorius Vivacon 500 #VN01H02) using 30 µg of protein resulting from each fraction. Samples were reduced, alkylated, and digested with trypsin (1:100) at 37°C for 14 hrs. Peptides were recovered from the filter using successive washes of 50 mM ammonium bicarbonate and 0.1% formic acid.

#### Cross-linked Amino Acid Analysis

Samples were processed using acid hydrolysis and liquid chromatography-tandem mass spectrometry (LC-MS/MS) analysis as previously reported^39^. Briefly, 5 mg of sample was placed in a glass vial for reduction with NaBH_4_ followed by acid hydrolysis over 24 hr using 6N HCl. Cross-links were enriched using flash cellulose chromatography before analysis via LC-MS/MS. Peak assignment and quantification were performed using MAVEN software and external standards were used for retention time assignments.

### Offline Fractionation

#### Size Exclusion Fractionation

Size exclusion fractionation was performed using an ÄKTA microsystem as previously described^40^. Peptides were separated on a Superdex Peptide PC 3.2/30 column (300 × 3.2 mm) at a flow rate of 50 μl min^−1^ using the SEC mobile phase (20% acetonitrile (ACN)). The separation was monitored by UV absorption at 214 and 280 nm. Two-minute fractions (100 μL) were collected into 96-well plates over a separation window of one column volume (∼2.4 mL). Fractions 1-3, representing high molecular weight peptides, were collected, evaporated to near dryness (Figure 4A), and analyzed by LC-MS/MS as described below.

#### High-pH Reverse Phase (HPRP) Fractionation

HPRP fractionation was performed using a Dionex 3000RS system as previously described^36^. Twenty-four fractions were created by combining fraction pairs evenly spaced across the separation gradient from 48 original fractions (i.e. fraction 1 with 25, fraction 2 with 26, etc.). The samples were partially dried and used for further analysis.

#### Strong Cation Exchange (SCX) Fractionation

SCX fractionation was performed using a Dionex 3000RS system as previously described^36^. Ten fractions were concentrated and cleaned using Pierce^TM^ C18 Spin Tips (Thermo Scientific #84850) according to the manufacturer’s protocol before further analysis.

### MS Acquisition

Global proteomic analysis was carried out on a Q-Exactive HF Orbitrap mass spectrometer (Thermo Fisher Scientific) coupled to an EASY-nLC 1200 (Thermo Fisher Scientific) through a nanoelectrospray LC-MS interface. Approximately seven μL of digested peptides (∼10 μg) were loaded on a fused silica capillary column (100 μm i.d. × 150 mm) packed in-house with 2.7 μm CORTECS C_18_ resin (Waters; Milford, MA). LC mobile phase solvents consisted of 0.1% FA in water (Buffer A) and 0.1% FA in 100% ACN (Buffer B, Optima™ LC/MS, Fisher Scientific, Pittsburgh, PA). After 22 μL of sample loading at a maximum column pressure of 700 bar, each sample was separated on a 120-min gradient at a constant flow rate of 400 nL/min. The separation gradient for cell fractions consisted of 4% buffer B from 0 to 3 minutes, followed by a linear gradient from 4 to 32% buffer B from 3 to 105 minutes. Gradient elution was followed by a linear increase to 55% buffer B from 105 to 110 minutes and further to 95% buffer B from 110 to 111 minutes. Flow at 95% buffer B was maintained from 111 minutes to 120 minutes to elute any remaining peptides. Data acquisition was performed using Xcalibur™ (version 4.5). The mass spectrometer operated in the positive ion mode. Each survey scan of m/z 300–1800 was followed by higher energy collisional dissociation (HCD) performed at a normalized collision energy of 30% collision energy, (Orbitrap Velos for HCD versus electron-transfer dissociation (ETD) comparison) with an isolation width of 1.6 m/z. The orbitrap was used for MS1 and MS2 detection at resolutions of 120,000 and 50,000, respectively. Dynamic exclusion was performed after fragmenting a precursor 1 time for 20.0 sec. Singly and doubly charged ions were excluded from HCD/ETD selection. Further details regarding the specific datasets can be found in supplemental table 3.

### Data Processing

#### Linear Peptide Searching and Quantification

Raw spectra were searched against a single database consisting of *Bos taurus* protein sequences (32,231 sequences, downloaded from UniProt on February 12, 2018) concatenated with mature collagen sequences using PEAKS X Pro. The precursor tolerance was set to ±15 ppm and fragment tolerance was set to ±0.08 Da. Fixed modifications were set as carbamidomethyl (C). Variable modifications were set as acetylation (protein N-term), deamidation (NQ), oxidation (M), oxidation (P) (hydroxyproline), and Gln -> pyro-Glu (N-term Q). The peptide and protein level false discovery rate (FDR) was 1%.

#### Initial identification of cross-linked peptides

Raw data was converted to peaklist using an in-house developed software (PAVA)^41^. Peptides were identified using Protein Prospector (v.6.3.1). A user-defined database containing the sequences of mature bovine COL1A1, COL1A2, COL2A1 and COL3A1 was used. Trypsin/hydroxylamine was specified as enzyme and a maximum of 1 missed cleavage site was allowed. Carbamidomethyl (C) and oxidation (M) were set as a fixed modification and variable modifications allowed were deamidation (NQ), oxidation (M), oxidation (P) (hydroxyproline), and Gln -> pyro-Glu (N-term Q). Deamidation (N) was defined as a “rare” modification; i.e. only one allowed per peptide. A user-defined cross-linker of H(-5)O(2)N(-1) between Lys and/or protein N termini was set to find HLKNL/deH-DHLNL-cross-linked peptides. A maximum of five variable modifications per peptide were allowed. Mass accuracies of ±10 and ±20 ppm were required for precursor and fragment ions, respectively and all *m/z* values were considered as monoisotopic values. The top 80 most abundant fragment ions were searched, and the instrument was defined as “ESI-Q-high-res”. Protein Prospector identifies cross-linked peptides using a mass modification searching strategy where cross-linked products are identified as linear peptides with undefined mass modifications on the site of cross-linking. In these searches it considered any mass modification between 283-6883 Da with a mass defect of 0.00048 per Da (to adjust for the typical mass defect from a peptide composition, allowing searches to be able to be performed with tight precursor mass tolerance) on lysine residues or the protein N-terminus. After performing this mass modification search the software looks for pairs of identifications to the spectrum that together add up to the mass of two linear peptides and the mass difference introduced by the cross-linker^42^. Acceptance parameters: minimum scores set to 22 and 15 and max E values set to 0.01 and 0.05 on the protein and peptide level, respectively. Minimum best discriminant score: 0, minimum MS- Tag low score: 0, maximum MS-Tag low E value: 1000, minimum score difference: 0.

To find additional cross-links, including trimeric products, the mass modification search results were inspected for spectra where one peptide was confidently identified (peptide score > 20), but the remaining mass was not explained. These were manually interpreted to try to explain the remaining mass.

#### Variable Modification Searching and Quantification

To identify additional cross-linked partners of peptides already confidently identified as members of cross-linked species with the above strategies, variable modifications were generated for identified cross-links by calculating the total mass for the most common modified forms of the COL1A1 and COL1A2 peptides of interest in combination with the mass alterations generated by the cross-link chemistry itself (Supplemental Table 2). Variable modifications were then employed by searching the main pulmonary artery data generated with the Q-Exactive HF mass spectrometer with PEAKS X Pro. The precursor tolerance was set to ±15 ppm and the fragment tolerance was set to ±0.08 Da. Data was searched against UniProt (32,231 sequences, downloaded 2/12/2018) restricted to *Bos taurus* with mature collagen sequences added. Fixed modifications were set as carbamidomethyl (C). Variable modifications were set as oxidation (M), oxidation (P) (hydroxyproline), and Gln->pyro-Glu (N-term Q) in addition to all custom cross-link modifications (Supplemental Table 2). Results were filtered to 1% FDR at the peptide and protein level. Peptides were also filtered if they contained an acetylation of an internal N-termini.

#### Spectral Library Generation

Spectral library generation (.MSP) was performed using an in-house Mascot server (Version 2.5, Matrix Science). The precursor tolerance was set to ±15 ppm and the fragment tolerance was set to ±0.08 Da. Data was searched against a custom database containing UniProt sequences for *Bos taurus* COL1A1 and COL1A2 with separate entries for the N- and C-terminal propeptides, as well as the mature collagen sequence. Additional proteins included in the database were bovine-specific COL2A1, COL6A3, and 16 ECM glycoproteins and proteoglycans in addition to 44 “decoy” sequences from other organisms to assess false positive cross-link identifications. Trypsin-specific cleavage was used in all searches. Fixed modifications were set as carbamidomethyl (C). Variable modifications were set as oxidation (M), oxidation (P) (hydroxyproline), Gln->pyro-Glu (N-term Q), α1NT_HLKNL (K), α1NT_LKNL (K), α2NT_HLKNL (K), and α2NT_HLKNL_Ox (K) (Supplemental Table 2). Results were filtered to 1% FDR at the peptide and protein level using Mascot. A spectral library containing 655 peptide sequences and corresponding spectral matches was generated using Mascot (Version 2.5).

#### Spectral Library Searching and Cross-link Quantification

Raw spectra for unfractionated bovine mPA were searched against the Mascot-generated spectral library using the MSPepSearch node within Proteome Discoverer (ThermoFisher, version 2.5). The precursor tolerance was set to ±10 ppm and fragment tolerance was set to ±0.08 Da. Results were filtered to a 1% FDR at the peptide level using the Fixed Value PSM Validator node and at the protein level using the Protein FDR Validator node. Label-free quantification (LFQ) was performed using the Precursor Ions Quantifier node.

## Results

Decellularized bovine ligament was utilized for the development of a method to detect peptides modified by LH and cross-linked by LOX, due to the high abundance of collagen I fibers present in this connective tissue. Two chemical digestion methods, hydroxylamine hydrochloride (HA) or cyanogen bromide (CNBr), were used to liberate protein from the bovine tendon samples. Following further trypsin digestion of the HA-digested sample, the resulting peptides were separated by strong cation exchange (SCX) or size exclusion (SEC) chromatography, and ten or three fractions, respectively, were isolated and analyzed by data-dependent LC-MS/MS (Figure 1E). A standard database search for linear peptides revealed that collagens I-III constituted greater than 99.75% (collagen I >99.45%) of the protein content of the previously insoluble starting material (Figure 1F). The overall coverage of the two collagen I chains was greater than 99% for the mature protein. We utilized amino acid analysis and identified several LOX-generated cross- links (DHLNL, HLNL, LNL, PYD, DPD) after reduction and complete acid hydrolysis of the protein. These results indicated that the di-hydroxylysine derived cross-link HLKNL/deH-DHLNL represents a significant percentage of collagen cross-links.

### Cross-linked N-telopeptide (NTX) identification

The first round of database searches to detect cross-linked products focused on HLKNL cross-links in only four collagen chains: COL1A1, COL1A2, COL2A1, and COL3A1. The database sequences for these fibrillar collagens include the N- and C-terminal propeptide sequences, limiting database recognition of this peptide with trypsin cleavage definitions. Therefore, we generated a custom database containing the sequences of the mature proteins, which allowed for the identification of the NTX peptides by database searching without the use of semi-specific or non-specific cleavage searches. The dominant cross-linked peptides identified after HA cleavage then tryptic digestion resulted from the cross-linking of the mature COL1A1 N-terminal peptide (α1NT, residues 1-25 of the mature protein; residues 162-186 of the database sequence) and COL1A2 N-terminal peptide (α2NT, 1-18 of the mature protein; residues 80-97 of the database entry) at residues K5 and K9 of the mature sequences (K170 and K84), respectively, via a mass addition that is consistent with a hydroxylysino-5-ketonorleucine (HLKNL) cross-link (Figure 2). The nomenclature used for these cross-links herein will be as follows: α1NT- α2NT_HLKNL. Identification of this cross-linked peptide was made using HCD and ETD fragmentation techniques (Figure 2A, B). Acquisition of ETD fragmentation spectra of the tryptic peptide yielded abundant c and z ions with excellent backbone fragmentation coverage (Figure 2B). However, ETD does not cleave N-terminal to prolines (or hydroxyprolines), and both peptides contain multiples of these residues. It should also be noted that ETD data acquisition was much slower (2.5 times more spectra were acquired in the HCD analyses compared to the ETD). As a result, a less comprehensive characterization of the sample was achieved using this mechanism of peptide fragmentation.

**Figure 2.**
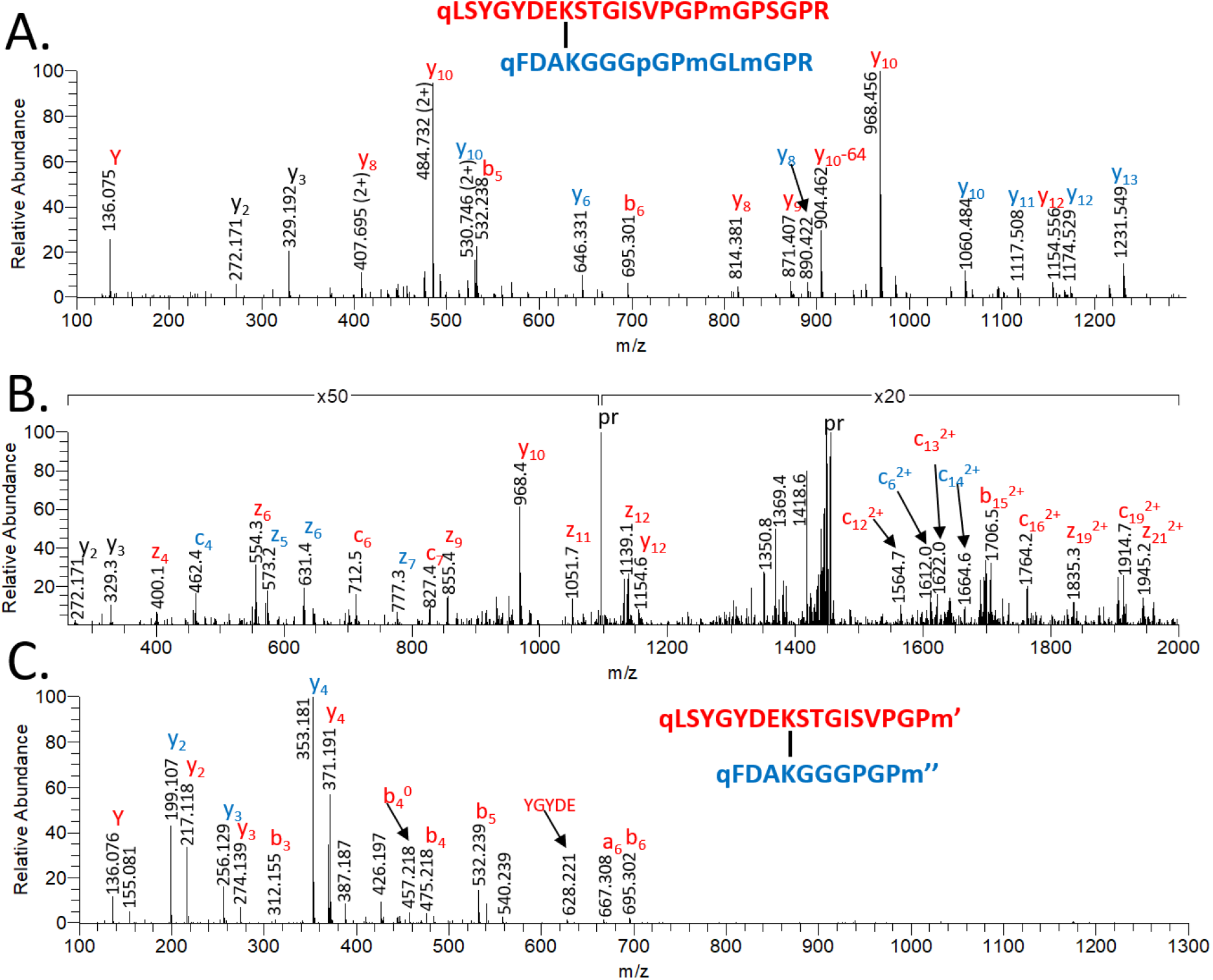
MS/MS spectra of the COL1A1-COL1A2 NTX dimer peptides cross-linked by the HLKNL linkage. COL1A1-derived fragment ions are highlighted in red, and COL1A2-derived fragments are in blue. Fragment ions in black can be assigned to both sequences in the cross-linked peptide. Low-intensity peaks corresponding to additional fragment ions are not labeled here. Modified residues: N-terminal pyroglutamic acid (q), oxidized Met (m), Hyp (p), homoserine (m’), and homoserine lactone (m’’). Fragment modifications: neutral loss of 64 Da (-64) and water loss (b_0_). A) HCD spectrum of precursor (pr) at m/z 1095.495 (4+). B) ETD spectra of m/z 1095.495(4+), representing the NTX dimer identified from the hydroxylamine treated sample. C) HCD spectrum of m/z 1030.800(3+), representing the NTX dimer identified from the CNBr treated sample. HCD spectra were acquired in the Orbitrap analyzer, and ETD spectra in the linear ion trap.

We also prepared samples using CNBr digestion, which we previously evaluated to access the chaotrope insoluble fractions from several tissues^37^. CNBr digestion generates peptides corresponding to 1-19 and 1-12 of mature α1NT and α2NT, respectively. Both of these peptides contain no basic residues other than the cross-linked residues that lose their charge when they are linked, and as a result, a less extensive series of fragment ions was observed compared to the hydroxylamine-pretreated samples (Figure 2C). For example, in the most fragment-rich spectrum of this cross-linked species, y ions for half the α2NT sequence were observed (6/12), whereas in the tryptic version derived from the hydroxylamine cleavage, 12/18 y ions were observed, providing a more confident identification (Figure 2A). Another disadvantage of this method is that heterogeneity is introduced as the terminal methionine residues are converted to homoserine and homoserine lactone species. Hence, we used HA cleavage for the remaining experiments.

### Trifunctional cross-link identification

There is currently no software for the identification of three peptides cross-linked through one trifunctional cross-linker from fragmentation data. The mass in a cross-linked product with an unexplained mass addition at the site of cross-linking in the case of a divalent cross-link is the mass of the second peptide and the elemental change due to the cross-linking reaction^42,43^. Based on confident identification of divalent cross-links between the two N-terminal peptides a similar approach was attempted with mass modification searching. The results were used to compare unexplained mass differences to the mass of the identified divalent species, to find a third peptide based on the extra mass. This approach was used to identify trivalent pyridinoline NTX peptides from the α1NT and α2NT peptides to the COL1A1 C-terminal helix region (α1CH, residues K946, K1107 UniProt entry) to give rise to α1NT-α2NT-α1CH_PYD (Figure 3A, B) and α1NT-α1NT- α1CH_PYD peptides (Figure 3C). These peptides eluted in the high-salt SCX fractions as expected based on containing three C-terminal Arg residues and at least one free N-terminal primary amine.

**Figure 3.**
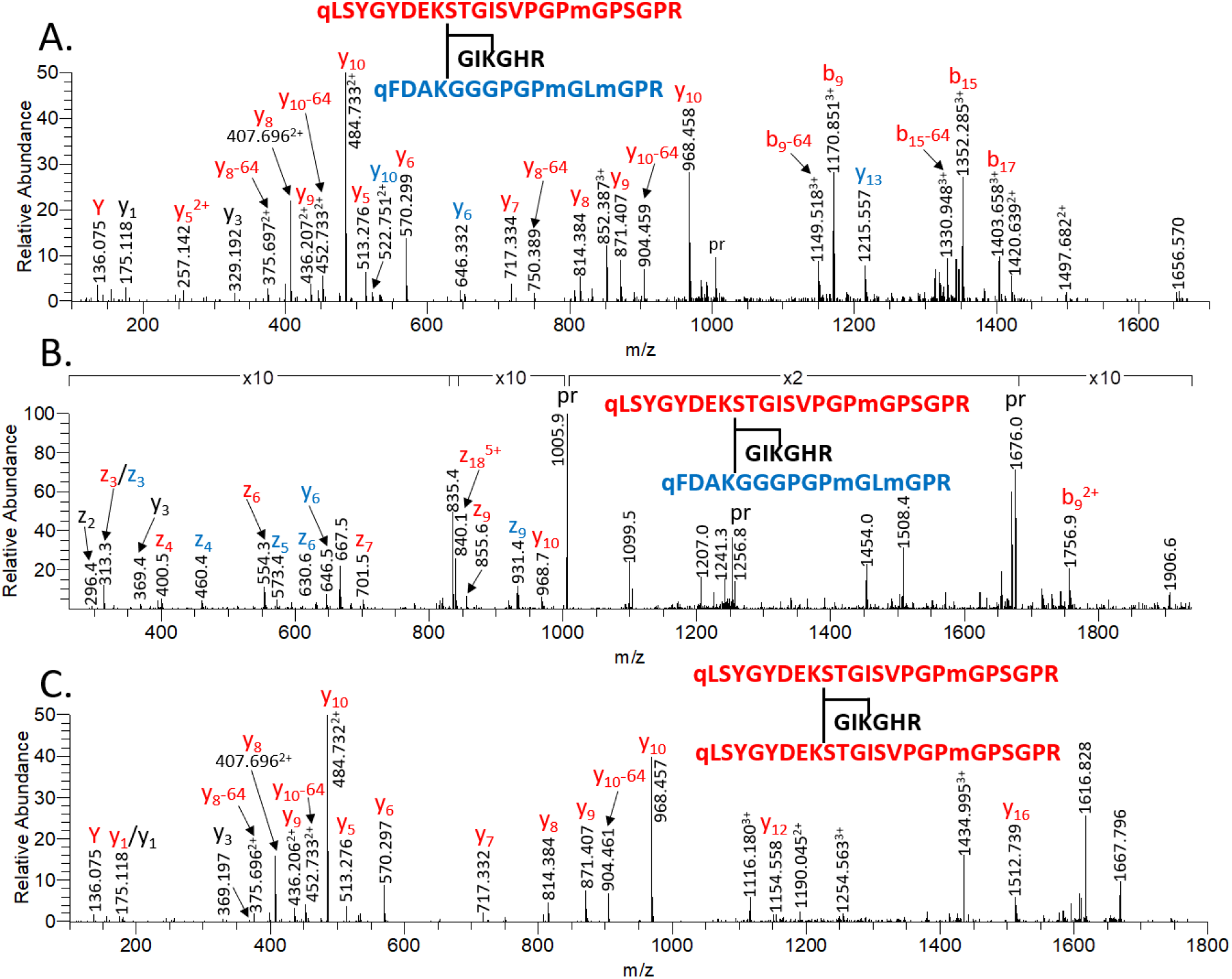
Trifunctional cross-linked NTX peptides with the COL1A1CH peptide. A) HCD spectrum of m/z 1005.263(5+) representing the α1NT-α2NT-α1CH_PYD trimer peptide identified via HA digestion. PYD = hydroxylysyl-pyridinoline linkage. The α1NT-related fragment ions are highlighted in red, α2NT-related fragments are in blue. Fragment ions in black can be assigned to both sequences. No specific fragment ions for the helical COL1A1 sequence were detected, its presence was only confirmed by the additive mass of the peptide. Modifications: N-terminal pyroglutamic acid (q), oxidized Met (m), neutral loss of 64 Da from fragment ion (-64, oxidized methionine side-chain loss). HCD spectra acquired in the Orbitrap analyzer (A & C). B) ETD spectrum of m/z 1005.263(5+) representing the α1NT-α2NT-α1CH peptide cross-linked by the PYD linkage identified in an HA-treated sample. α1NT-derived fragment ions are highlighted in red, α2NT-derived fragments are in blue, and α1CH-derived fragments are in black. ETD spectrum acquired in the linear ion trap. C) HCD spectrum of m/z 1163.544(5+) representing the α1NT- α1NT-α1CH trimer peptide cross-linked by the PYD linkage identified in an HA-treated sample. α1NT-derived fragment ions are highlighted in red, α1CH-derived fragments are in black.

### N-Telopeptide Heterogeneity

The linear N-telopeptides of mature collagen I were identified with and without one trypsin missed cleavage. Analysis of common PTMs revealed a wide range of identifications for the linear (non-cross-linked) tryptic COL1A1 N-telopeptide (α1NT, residues 1-28). The first N-terminal peptide (residues 1-9) was not identified, but the second tryptic peptide (residues 10-28) was identified with one, two, and three oxidation/hydroxylations (M, P, and K). These peptides account for almost 87% of the total area under the curve (AUC) signal for the linear α1NT sequence. The majority of the missed-cleaved form (residues 1-28) was found with the N-terminal glutamine in the pyro-Glu cyclic state (∼12.5% α1NT AUC signal) across 1-4 oxidations, with 2 accounting for the largest contribution to signal (7.8%). Minor forms were identified from four alternative states for the N-terminal Q; unmodified (0.2%), deamidated (d, 0.5%), acetylated (Ac, 0.3%) and acetylated and deamidated (Ac-d, < 0.1%). Of note, several of these minor component identifications are not included within a 1% FDR cutoff. The acetylation of mature telopeptides is surprising as N-terminal acetyltransferases (NATs) are expected to function at the ribosome or golgi, locations at which the propeptide regions of collagen should still be present. Acetylation is also not a likely side reaction of sample preparation under the conditions used.

### Variable modification candidate cross-link identification

From the variable modification searches a total of 374 unique NTX candidate cross-linked peptide spectra were identified across 24 proteins at an estimated 1% FDR. Most of these candidates are core ECM components (91%), collagens (90%), fibrillar collagens (85%), and specifically collagen I chains (81%). These identifications were filtered to include only those that were observed in two or more of the analyzed datasets. This narrowed the candidates down to 68 cross-linked species. All were on the N-terminal peptide of COL1A1 with variations in the number and site/s of oxidation/hydroxylation and N-terminal glutamine status. The most common modifications were hydroxylation of P16 and P18 and oxidation of M19. N-terminal glutamine states varied between unmodified, deamidated and pyro-glutamate.

Table 3 shows the unique cross-link candidates for the NTX COL1A1 peptides by dataset (identifications are grouped by cross-link mass addition). The total number of candidates can be found in Supplemental Materials. For the tendon samples, several of the analytical combinations gave rise to a high number of unique cross-link species. For example, HA digestion with SCX (set B) gave the highest number of unique candidates, whereas the use of SEC yielded the most complete variations of the NTX candidates, with identifications in all XL candidate categories. Per LC-MS/MS run, the NTX identifications were highest for the SEC dataset, however, the unique cross-link identifications per MS2 scan were highest for the SCX dataset acquired at the same time at ∼6/10,000 scans, versus ∼5/10,000 scans for the SEC dataset.

**TABLE 3.**
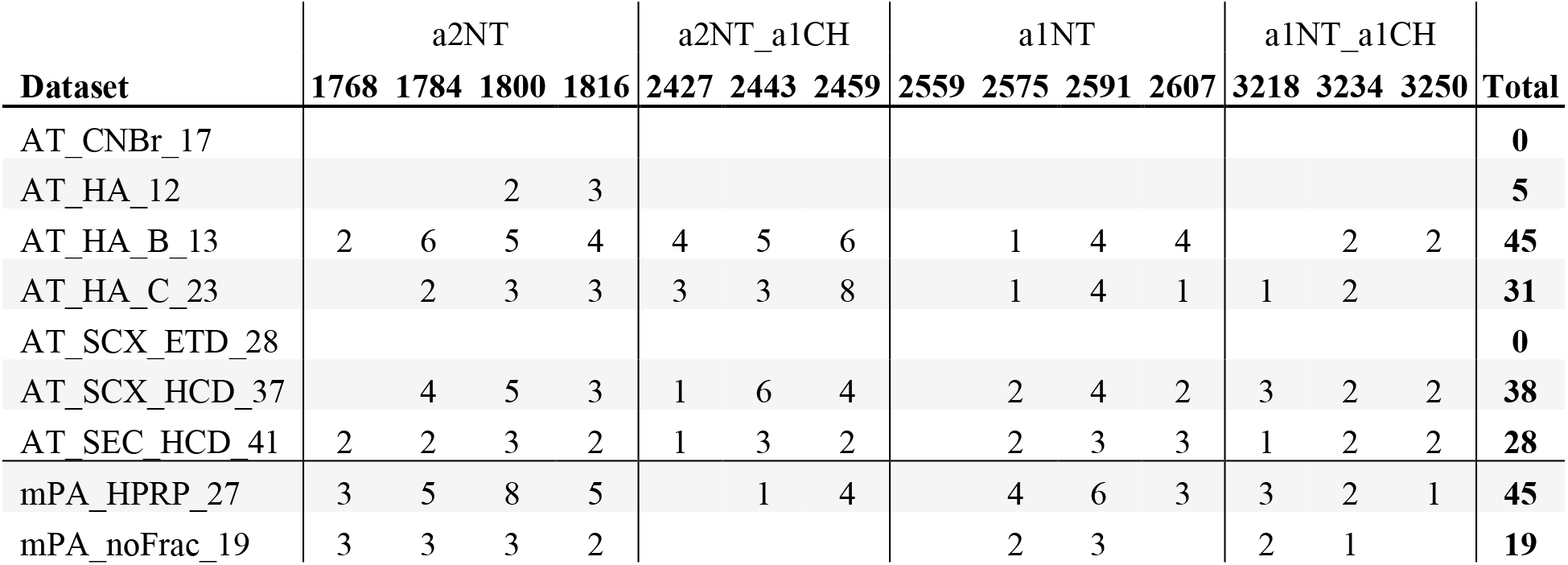
Unique collagen I N-terminal telopeptide identifications using a variable modification search strategy. The specific variable modifications are shown based on the integer of the exact mass. For example, 1799.7607 is COL1a2NT_HLKNL (See Supp. Table 2). Tissue: Achilles tendon (AT), main pulmonary artery (mPA). Digestion: Cyanogen Bromide (CNBr), Hydroxylamine (HA, if not specified then HA digestion was used). Fractionation: strong cation exchange (SCX, if not specified SCX was used), size exclusion chromatography (SEC), high pH reversed phase (HPRP), no fractionation (noFrac).

The extent of diversity in PTM states observed in both linear and cross-linked collagen peptides is especially evident when analyzing raw LC-MS/MS spectra (Figure 4). Analysis of cross-link containing SEC fractions yields peptide ion envelope trains with separation occurring on both the retention time (RT) and mass-to-charge (m/z) axes. The separation observed on the m/z axis is largely driven by mass additions with increasing numbers of hydroxyprolines and hydroxylysines. Separation on the RT axis is driven by PTM positional isomers and, in some cases, possible proline conformation.

**Figure 4.**
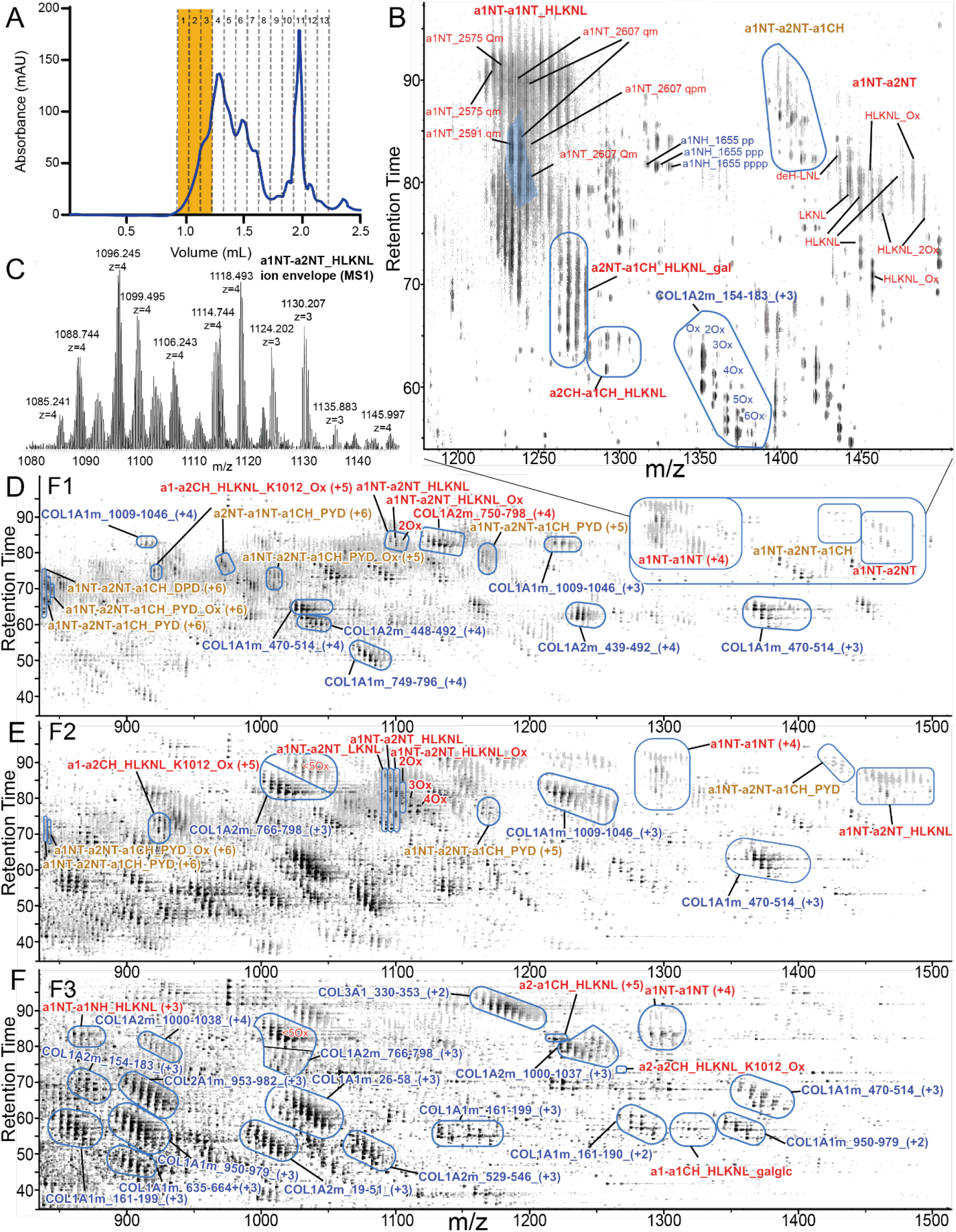
LC-MS spectra highlighting the ion envelope landscape of large linear and cross- linked collagen peptides. SEC fractionated bovine tendon MS1 spectra with identified peptides annotated. A) Chromatogram trace of SEC fractionation, with the fractions analyzed highlighted in yellow. B) Enlarged panel highlighting a region within F1. The shaded blue region represents the total LFQ-quantified area for α1NT-α1NT_HLKNL. The quantified proportion of the ion envelope is consistent across most cross-linked peptide identifications. C) MS1 trace of the ion envelope for α1NT-α2NT_HLKNL seen in F2. F1-3 represents the first three fractions collected. The regions outlined in blue represent the ion envelope for the annotated identification. D) F1 contains the most abundant trifunctional cross-linked collagen peptide identifications. E) F2 contains mostly difunctional cross-linked peptides. F) F3 consists mostly of linear collagen peptides. Modifications: N-terminal pyroglutamic acid (q), oxidized Met (m), Hyp (p).

Despite accounting for an extensive list of PTM states and cross-linked peptide arrangements (Tables 1 & 2), LFQ values underrepresent the abundance of high m/z collagen peptides in most cases. This is especially evident in modified cross-linked peptides, as highlighted in Figure 4B. The region of the α1NT-α1NT_HLKNL ion envelope shaded in blue represents the total quantified area. This region contains the highest intensity peaks in the ion envelope but represents less than 1/3 of the total area. This is consistent across most cross-linked collagen peptide ion envelopes as well as some heavily modified linear peptide ion envelopes, with only the most common isoforms being selected for quantification.

### NTX Cross-link Analysis in Disease

To test if NTX peptides could be identified in tissue samples with more diverse proteomes, we utilized a compartment-resolved proteomics approach^44^ to generate cellular and ECM fractions from main pulmonary artery (mPA) samples in a bovine hypoxia-induced model of pulmonary hypertension (hPH)^45,46^. In 1915, a syndrome of right heart failure was identified in cattle raised at high altitudes in Colorado^47^. Later studies demonstrated that exposure to low levels of oxygen induces alveolar hypoxia, consistently resulting in chronic pulmonary hypertension and alterations to the structure of precapillary pulmonary vessels that recapitulate human disease better than rodent models^45,48^. hPH generates arterial remodeling involving fibroblast activation with collagen deposition and increased biomechanical stiffening of the vessels^49^. In this study, tissue hydrolysates revealed significant increases in cross-linked amino acids in tissues from animals with PH versus controls (Figure 5A). The most significant increase was found in the LH and LOX-mediated cross-link deH-LNL (4.6-fold increase) At the proteomics level, we observed increased levels of cross-linking-associated enzymes LOX, LOX-like 1 (LOXL1), and lysine hydroxylase 2 (PLOD2/LH2) (Figure 5A).

**Figure 5.**
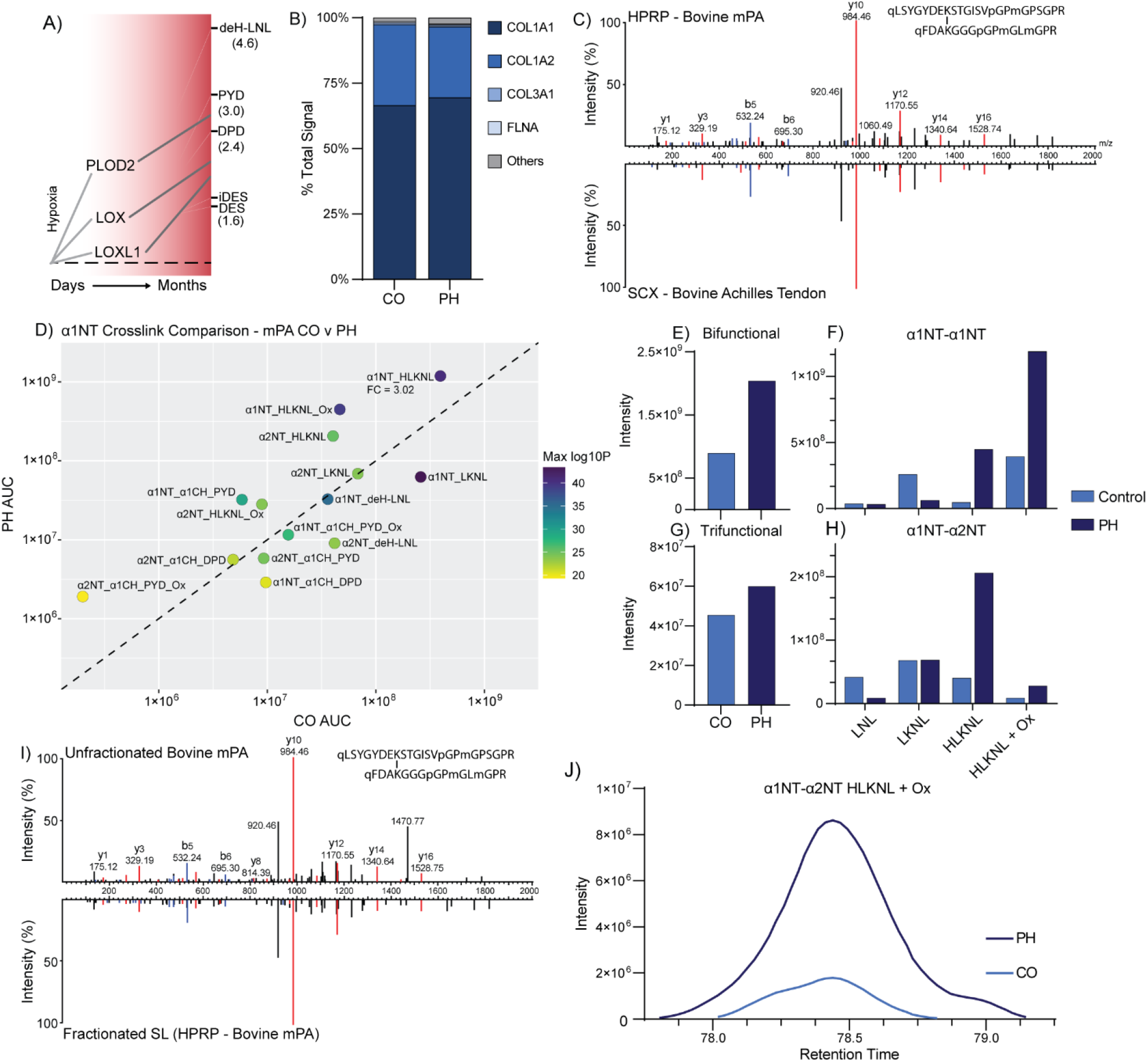
Variable modification searching for identification and quantification of NTX cross- links in bovine pulmonary hypertension. A) Summary of protein and cross-link changes throughout pulmonary hypertension progression. Fold changes for cross-links are displayed in parentheses. B) Proteomic composition of insoluble ECM fractions from control (CO) and pulmonary hypertension (PH) main pulmonary artery (mPA) by area under the curve intensity. C) α1NT-α2NT_HLKNL spectra from SCX fractionated bovine Achilles tendon (bottom) and HPRP fractionated bovine mPA (top). D) Comparison by intensity of all α1NT cross-links identified in both mPA groups via variable modification searching between CO and PH mPA. Fold change is calculated as PH/CO and is displayed for cross-links including the α1NT peptide. Points are colored by the maximum log10 p-value for the identification of each species. E-H) Comparison of identified intensity for bifunctional (E), trifunctional (G), and all cross-links by oxidation for α1NT-α1NT (F) and α1NT- α2NT (H) between CO and PH mPA. I) Spectrum match for α1NT- α2NT_HLKNL_Ox identification in unfractionated bovine mPA using a spectral library searching approach. A spectral library was generated using the HPRP fractionated bovine mPA data displayed in (C). J) Extracted ion chromatograms for identification of α1NT-α2NT_HLKNL_Ox in unfractionated CO and PH mPA (m/z 1465.66, 3+).

We utilized HA and trypsin digestion followed by offline chromatography to generate 10 fractions from disease and healthy control mPA tissue. Consistent with the cross-linked amino acid analysis and increased cross-linking enzymes, several NTX species were identified and found to increase with hPH, including all HLKNL bifunctional cross-links α1NT-α1NT_HLKNL and α1NT-α2NT_HLKNL, as well as prominent trifunctional cross-links α1NT-α1NT-α1CH_PYD and α1NT-α2NT-α1CH_PYD (Figure 5D). However, some cross-links, including α1NT- α1NT_LKNL, α1NT-α2NT_deH-LNL, and α1NT-α1NT-α1CH_PYD decreased with PH, indicating that rearrangement of cross-link types/specificity occurs alongside an overall increase in cross-link abundance during hPH. Overall, a 2.3-fold increase in abundance was observed for bifunctional while a 1.3-fold increase was observed for trifunctional cross-links, consistent with new/immature collagen cross-links being formed in a lysyl hydroxylase-dependent pathway with disease progression (Figures 5E and 5G).

Varying changes in cross-linked peptides were observed depending on the number of oxidative modifications (Figure 5F, H). For the α1NT-α1NT cross-link, the deH-LNL and LKNL linked species abundance did not change and decreased by 4-fold with PH, respectively, while HLKNL and HLKNL + Ox abundance increased by 9.6- and 3-fold, respectively, to become the species with the most intense signal (Figure 5F). Similar trends were observed for the α1NT-α2NT cross-link, where deH-LNL and LKNL showed no increase with PH while HLKNL and HLKNL + Ox forms increased by 5.1-fold and 3.2-fold, respectively (Figure 5H). Again, this is consistent with proteomics-level data showing increases in PLOD/LH enzymes responsible for the oxidation of lysine residues before LOX cross-linking.

To determine if some of these NTX species could be identified without the offline fractionation involved in the discovery phase of experiments, mPA samples from cattle that developed PH versus those that did not were evaluated. These samples were enriched for the ECM but not fractionated at the peptide level and acquisition was not biased towards higher charge state peptides. A spectral library for a subset of the NTX peptides identified in the fractionated data was used to search this unfractionated data. A total of 5 NTX species in different modification states were identified at the FDR cutoff across 54 total spectral matches. The spectra assigned to α1NT- α2NT_HLKNL_Ox were readily matched to the spectral library entry (Figure 5I). Quantification by AUC indicates that this species of NTX peptide was approximately 8-fold higher in PH than in CO samples (Figure 5J).

## Discussion

Recent advancements in chemical cross-linking reagents, data acquisition methods, and data analysis routines have positioned cross-linking mass spectrometry (XL-MS) as a prominent tool for the structural study of proteins. Many of these methods can also be used to explore cross- links generated *in vivo*. For example, factor XIIIa-generated cross-links have been mapped in blood clots and clinical thrombi using this approach^36^. Identification and mapping of the cross- links found in the ECM can yield valuable information about disease-generated alterations to the architecture of tissues and resulting mechanical forces exerted on embedded cells. For example, we and others have shown drastic architectural changes in collagen fibers as revealed by second harmonic generation (SHG) imaging in tumors, pulmonary hypertension, and other fibrotic conditions in which fiber density and alignment characteristics correlate with disease progression^50,51^.

Challenges associated with LOX cross-linked peptide identification include the parent proteins residing in a tissue fraction that is resistant to detergent extraction, proteolytic cleavage sites that are protected due to tight fiber bundling, heterogeneity at the peptide and cross-linked amino acid levels, and large mass/charge states in the gas phase. Some of these features can also allow for targeting of these PTM-derived products. For example, the hydrophobicity of cross- linked species can be utilized for enrichment, and high *m/z* values and/or precursor charge states can be targeted for data acquisition and identification.

In addition, there are multiple layers of variability in collagen I N-telopeptides. First, there are multiple modification states based primarily on N-terminal glutamine modification with up to five states, one to four additions of oxygen on the α1NT peptide (M, P and K), and deamidation, in part due to the long half-life of collagen *in vivo*. The second level of complexity arises from the combinations of lysine PTMs associated with the cross-link products. Evidence for an additional oxidation of the di-hydroxylysine-derived cross-links is also supported by our findings (data not shown). The addition of O-linked galactosyl and galactosylglucose must be considered when considering the C-helical sites as well. Based on these variables, there are well over a hundred possible combinations of the NTX peptide. This heterogeneity further increases at the level of the cross-link state which can reside as aldimines and ketoamines when derived from hydroxylysine^52^ along with the reduced forms. Evidence for more than 50 of these has been presented here.

Here, we utilized HA or CNBr to cleave the peptide backbone and liberate cross-linked peptides from mature collagen fibers. The HA reaction exploits cyclic imide formation of asparagine residues at sites N228 and N180 in the COL1A1 and COL1A2 N-telo regions, respectively^53^. Unlike enzymatic cleavage, these reactions should be much less sterically hindered by fiber packing. HA digestion is accelerated by the strong chaotrope/denaturant guanidine hydrochloride to yield protein fragments that are amenable to further enzymatic digestion^37^. While the HA approach used here is effective at digesting insoluble protein and has the advantage of pushing Met oxidation to completion, it may induce additional oxidative modifications^44^. In general, HA cleavage resulted in many more collagen I NTX identifications than using CNBr, and spectra of greater quality due to improved fragmentation properties from the presence of the basic C-terminal residues. The peptides from both digests produced few N-terminal fragment ions due to the lack of the primary amine at the N-termini, with pyro-Glu formation from N-terminal Gln of both proteins representing the most common species.

The N-terminal cross-linked species found in collagen I fibers were characterized, and the resulting spectra revealed a wide array of unique peptide species arising from a diverse set of additional PTMs. Taking these modifications into consideration creates a large database search space even when a small subset proteome database is used. Collagen presents a particular challenge based on the presence of GPP repeats, the number of enzymatically catalyzed PTMs observed, and the heterogeneous nature of those modifications (hydroxyproline, hydroxylysine, glycosylation, deamidation, advanced glycation end products, pyro-Glu formation, etc.). An additional confounding factor is that for large *m/z* species, the monoisotopic peak of a peptide ion envelope is often not detected, and the resulting peptide spectra are associated with an incorrect monoisotopic *m/z*. This can be accounted for by using a high-resolution instrument and a modern database search algorithm allowing for selection of the +13C peak or even +13C(2) peak correction. When considering cross-linked species that only differ by two hydrogens, such as HLKNL/deH-DHLNL and the reduced form DHLNL, an incorrect cross-linker assignment can occur.

While 77 unique cross-linked species were identified, most of the heterogeneity originated from the cross-link species, hydroxyproline, oxidized methionine and N-terminal glutamine status. Some of the unique species are from positional isomers of P-OH. This is likely slightly overestimated due to challenges with the identification of PTM positional isomers and the real possibility that some of the unique positional isomers are mis-assigned. However, the NTX peptide has a minimum of several dozen forms in the two tissues studied here.

Fiber diffraction has been used to explore the atomic structure of collagen fibers for almost a century^54^. The diffraction data presented by Orgel et. al. revealed new features of collagen secondary structure, fiber pack, and overall quaternary orientation^55^. Several intra- and inter-fiber densities were not explained by the model (Figure 6A). The abundant α1NT-α2NT-α1CH_PYD identified cross-link fits well within the observed electron density from the previously reported model of the rat tail collagen structure (Figure 6B) providing support for the trifunctional cross- linked species identified here.

**Figure 6.**
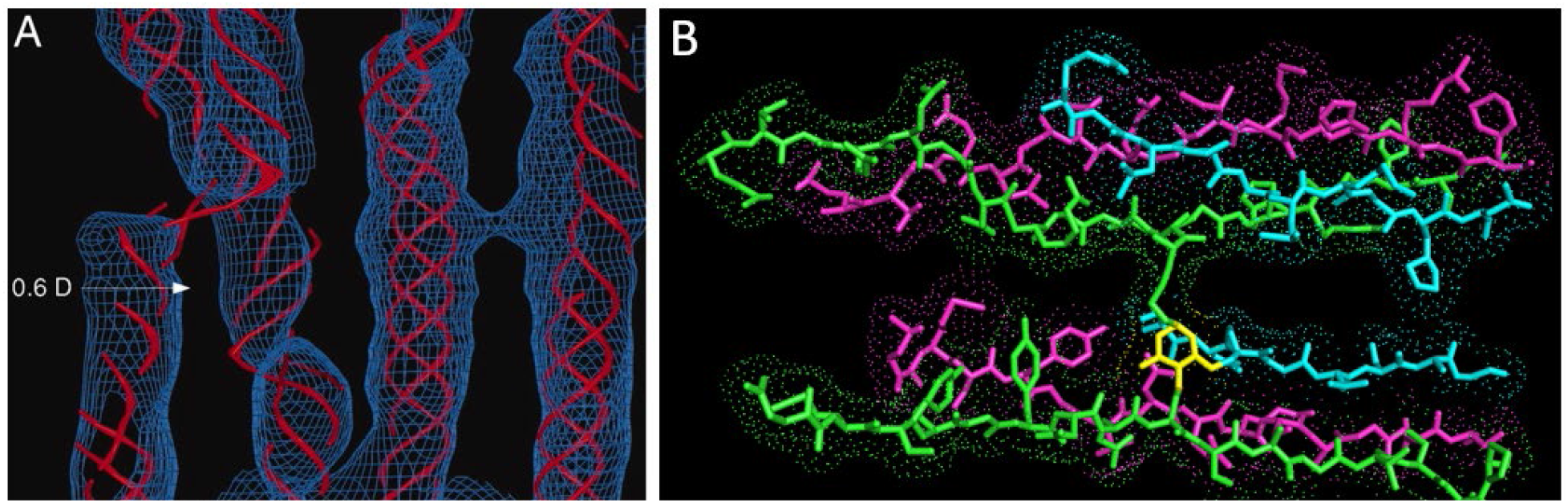
X-ray diffraction reveals unassigned density in position of collagen cross-links. A) Density map generated by x-ray fiber diffraction displaying the region of overlaid COL1A1 and COL1A2 chains containing the NTX peptide^52^. B) α1NT-α2NT-α1CH_PYD cross-link modeled into missing density between COL1A1 and COL1A2 peptides.

Once identified, the resulting assigned spectra can be used in spectral library search methods to facilitate more routine analysis. Using this approach, a subset of the NTX peptides identified in a discovery experiment with offline fractionation were quantified in mPA samples from a bovine model of hypoxia-induced pulmonary hypertension. Identification of the NTX peptides and their increase with hPH is consistent with increased levels of LH and LOX protein levels and resulting cross-linked amino acid products of these enzymes.

Despite the use of high-resolution MS with an accurate mass dataset (3σ < +/-2 ppm), a large percentage of search results are false positives. This finding highlights the importance of evaluating score differences to the next best peptide matches and independent identification of the two peptides from fragmentation spectra^42^. False positive identifications will be minimized by utilizing proteases with high specificity and high-quality tandem data that yield confident matches for the peptides in a cross-link. Identification from orthogonal digestion approaches and additional data that is consistent with the candidate identification (i.e. chromatographic elution) provide additional support. Generation of the cross-linked species using *in vitro* reconstitution is extremely challenging as PLOD and LOX systems are sparse due to difficulties with protein production and solubility. In addition, the synthesis of cross-links is challenging despite the development of orthogonal protection strategies required for incorporation into peptides. For these reasons, validation of cross-linked species with standards will remain challenging. Alternative digestion approaches, good coverage of fragment ions from both peptides, through cross-link fragments, and expected chromatographic elution behavior, can all be used to support a given assignment.

Structural studies of large ECM structures such as elastic or collagen fibers will remain challenging due to the flexible/non-crystalline nature and heterogeneity observed in these macromolecular complexes. However, the XL-MS approach presented here can yield information about protein-protein topology and how it changes with disease progression and aging. The findings can be used to improve 3-dimensional models, characterize the function of individual LH and LOX enzymes and aid in the evaluation of therapeutic interventions to block or reverse this component of matrix remodeling. The complexity observed at the N-telopeptide of collagen I here is reminiscent of the disordered region of histone N-termini/tails. The PTMs and resulting structures may create a “collagen code” that instructs cellular response/phenotype through mechanical and biochemical queues in tissue microenvironments in an analogous manner. The approaches outlined here can be used to evaluate such hypotheses.

## Author Contributions

The manuscript was written through contributions of all authors. / All authors have given approval to the final version of the manuscript. / #These authors contributed equally.

## Data Availability

The MS proteomic data have been deposited to the ProteomeXchange Consortium via the PRIDE partner repository with the dataset identifier PXD043595.

## Acknowledgements

This work was supported by the NIH (Grant Nos. R33CA183685, RM1GM131968, P01HL152961, and R01HL146519) and the University of Colorado Cancer Center Support Grant (P30CA046934), NIH grant P41GM103481, the Howard Hughes Medical Institute and the Dr Miriam and Sheldon G. Adelson Medical Research Foundation. We would like to thank Monika Dzieciatkowska and Mark A. Burlingame for maintaining the analytical platforms used for this work.

## Acronyms

COL: collagen
ECM: extracellular matrix
NC1: non-collagenous domain
LOX: lysyl oxidase
LH/PLOD: lysyl hydroxylase (gene: procollagen-lysine,2-oxoglutarate 5-dioxygenase)
LC-MS/MS: liquid chromatography tandem mass spectrometry
ETD: electron-transfer dissociation
HCD: higher energy collisional dissociation
NTX: mature collagen I N-terminal cross-linked peptide
CTX: mature collagen I C-terminal cross-linked peptide
LNL: lysinonorleucine
deH-LNL: dehydro-lysinonorleucine
HLNL: 5-hydroxylysylnorleucine
LKNL: lysino-5-ketonorleucine
deH-HLNL: dehydro-hydroxylysinonorleucine
DHLNL: dihydroxylysinonorleucine
HLKNL: hydroxylysino-5-ketonorleucine
deH-DHLNL: dehydro-dihydroxylysinonorleusine
PYD: pyridinoline
DPD: deoxypyridinoline
Hyl: hydroxylsine
Lys: lysine
alLys: allysine
alHyl: hydroxy-allysine
Arg: arginine
P-OH: hydroxyproline
CNBr: cyanogen bromide
mPA: main pulmonary artery
hPH: hypoxia-induced pulmonary hypertension
HA: hydroxylamine
SHG: second harmonic generation imaging
SCX: strong cation exchange chromatography
SEC: size exclusion chromatography
XL-MS: cross-link mass spectrometry
α1NT: collagen I alpha-1 mature N-terminal peptide
α2NT: collagen I alpha-2 mature N-terminal peptide
α1CH: collagen I alpha-1 C-helical peptide (cross-linking site)

## Supplemental Material

**Supplemental Figure 1.**
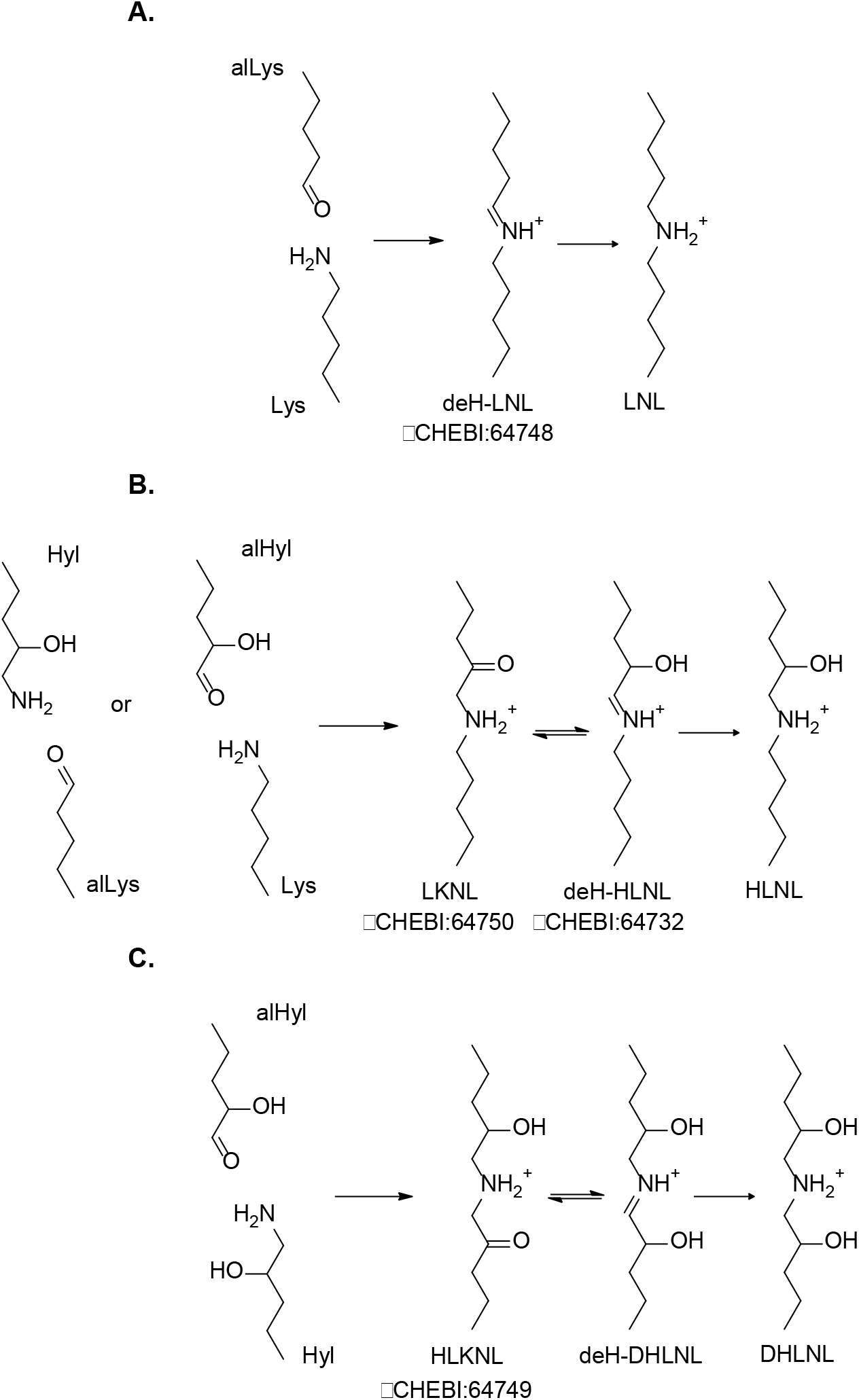
Difunctional cross-links. Formation of LNL from alLys and Lys (A), HLNL from Hyl and alLys or alHyl and Lys (B), and DHLNL from alHyl and Hyl (C).

**Supplemental Table 1.**
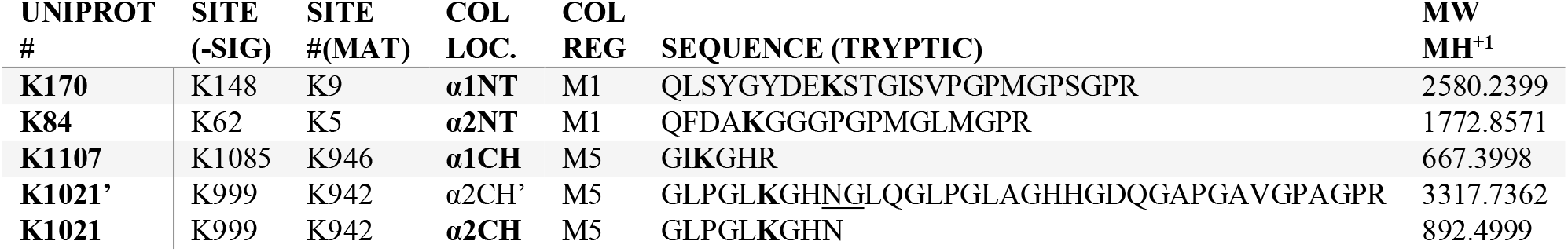
Collagen NTX Candidates. Tryptic peptides with Lysine residues in rough alignment with the N-terminal telopeptides based on the D-banding and fiber X-ray diffraction models. Reported NTX cross-link containing peptides in their full cleavage state are in bold. The mature sequences (MAT) start after the signal residues 1-22 for both 1a1 and 1a2 and propeptide residues 23-161 and 23-79 for 1a1 and 1a2 respectively. UniProt entries P02453 and P02465. Lysine cross-linking sites and hydroxylamine cleavage sites are in bold, low probability cleavage sites are underlined. K#’ denotes the hydroxylamine mis-cleaved form of the peptide.

**Supplemental Table 2.**
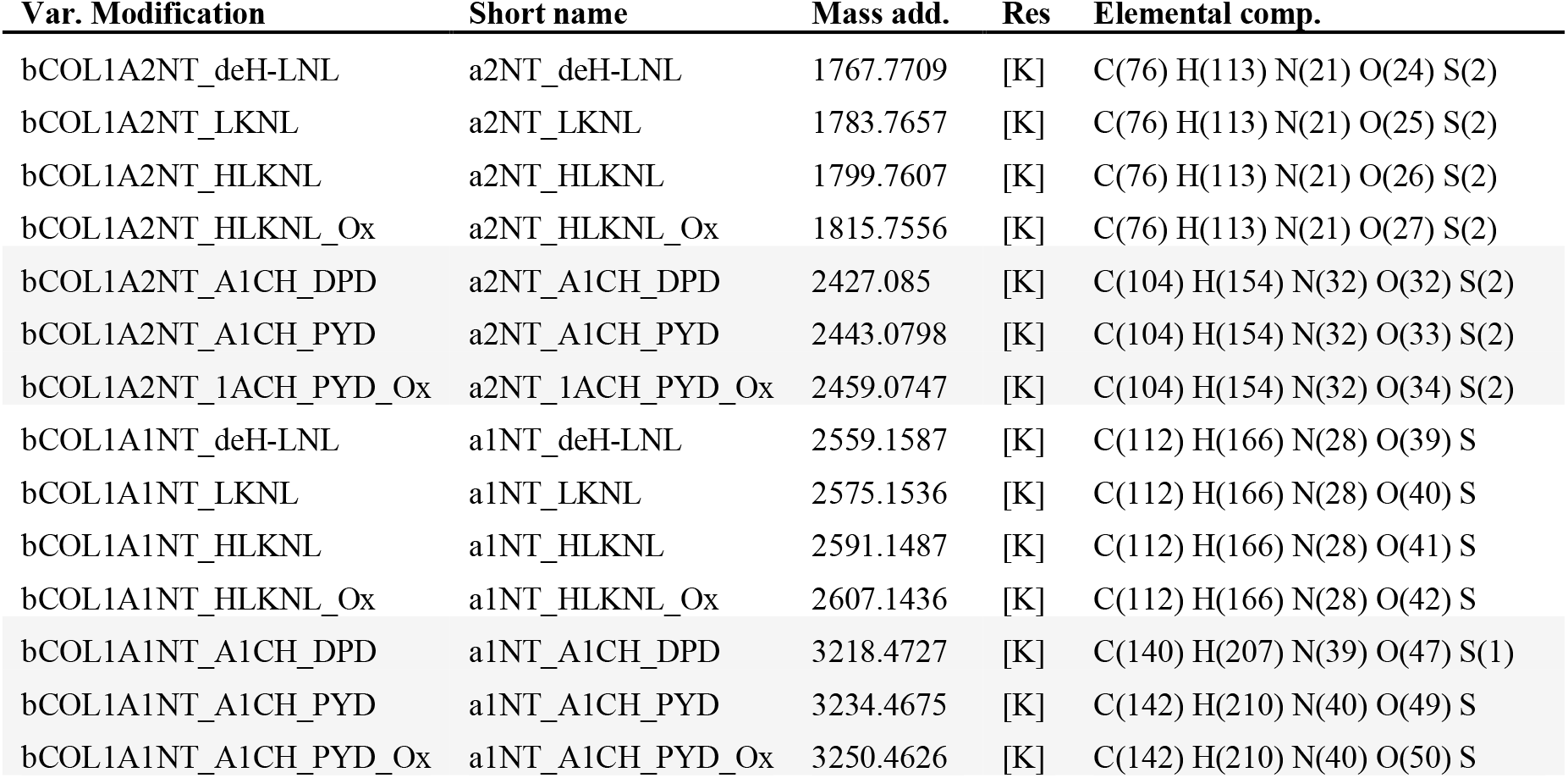
NTX variable modifications (PEAKS searches)

**Supplemental Table 3.**

| **Dataset Name\*** | **cleavage specificity** | **fract. method** | **MS Inst** | **Activ.** | **run length** | **Files / Frac.** | **MS1 scans** | **MS2 scans** | **#Features** |
| --- | --- | --- | --- | --- | --- | --- | --- | --- | --- |
| AT_CNBr_17 | Tryp/CNBr | SCX | QE HF | HCD | 90 | 10 | 147635 | 251958 | 223050 |
| AT_HA_12 | Tryp/HA | SCX | QE HF | HCD | 90 | 9 | 116281 | 243967 | 246856 |
| AT_HA_B_13 | Tryp/HA | SCX | QE HF | HCD | 90 | 10 | 23765 | 238144 | 162186 |
| AT_HA_C_23 | Tryp/HA | SCX | QE HF | HCD | 90 | 7 | 12261 | 183441 | 120957 |
| AT_SCX_ETD_28 | Tryp/HA | SCX | Orbitrap Velos | ETD | 78 | 10 | 28317 | 64100 | 205636 |
| AT_SCX_HCD_37 | Tryp/HA | SCX | Orbitrap Velos | HCD | 78 | 10 | 28317 | 64100 | 205636 |
| AT_SEC_HCD_41 | Tryp/HA | SEC | QE+ | HCD | 120 | 3 | 20489 | 63034 | 163628 |
| mPA_HPRP_27 | Tryp/HA | HPRP | Fusion Lumos | HCD | 70 | 24 | 158395 | 323323 | 2992546 |
| mPA_noFrac_19 | Tryp/HA | none | QE HF | HCD | 120 | 4 | 19668 | 294965 | 159917 |

\* names match samples and results found in the PRIDE entry PXD043595.

